# Monarchs sabotage milkweed to acquire toxins, not to disarm plant defence

**DOI:** 10.1101/2023.03.20.533455

**Authors:** Anja Betz, Robert Bischoff, Georg Petschenka

**Affiliations:** Department of Applied Entomology, University of Hohenheim, Stuttgart, Germany

## Abstract

Sabotaging milkweed by monarch caterpillars is a textbook example for disarming plant defence. By severing leaf veins, monarchs are thought to prevent toxic latex flow to their feeding site. Here we show that sabotaging by monarch caterpillars is not an avoidance strategy. Instead, caterpillars actively ingest outflowing latex to increase sequestration of toxic latex cardenolides. Comparisons with caterpillars of the related non-sequestering common crow butterfly revealed three lines of evidence supporting our hypothesis. First, monarchs sabotage inconsistently and therefore the behaviour is not mandatory to feed on milkweed, while sabotaging in crows precedes every feeding event. Second, monarchs eagerly drink latex, while crow caterpillars spit out latex during sabotaging. Third, monarchs raised on detached leaves sequestered more cardenolides when latex was supplemented artificially. Hence, we conclude, that monarchs converted the “sabotage to avoid” strategy of their relatives into a “sabotage to consume” strategy for acquiring toxins for defence.

## Main Text

Upon mechanical injury, roughly 9% of all angiosperm plants exude latex, which often contains toxins and functions as a chemical and physical barrier against insect herbivores by poisoning them or gumming up their mouthparts ^1–5^. Before taking up food, many plant-feeding insects therefore sabotage the latex-containing elongated cells (i.e. laticifers) running along the leaf veins and cut off leaf veins or petioles with their mandibles to drain the feeding site and to circumvent exposure to latex ^6,7^. This sabotaging behaviour has been observed in various, unrelated insects including caterpillars, beetles, and katydids ^8–10^ and likely represents a common response of insect species feeding on latex-bearing plants ^11^.

Sabotaging behaviour has been especially well studied in caterpillars of the monarch butterfly (*Danaus plexippus*) feeding on toxic milkweed (*Asclepias* spp.), and represents a textbook example for the disarming of a plant defence trait by an insect ^12–18^. Besides being sticky ^19^, latex of many milkweed species contains high concentrations of cardenolides ^20^, potent toxins inhibiting Na^+^/K^+^-ATPase, an enzyme essential for many physiological processes in animals ^21–23^. Remarkably, monarchs cannot only cope with cardenolides by means of a resistant Na^+^/K^+^-ATPase ^24^ but are also famous for sequestering cardenolides in their body tissues as a defence against predators ^25–27^.

When sabotaging plants, monarch caterpillars disable laticifers either by severing all minor laticifers supplying a small section on a leaf (“trenching”), or by cutting major veins which supply entire leaf portions with latex (“vein cutting”) (Fig. 1) ^11,28^. Notably, sabotaging behaviour is not restricted to *D. plexippus* but has also been observed in related milkweed butterflies including *Danaus erippus, Danaus gilippus, Euploea core, Euploea crameri, Idea leuconoe, Lycorea cleobaea*, and *Parantica sita* feeding on latex-containing plants of the Apocynaceae, Moraceae and Caricaceae ^3,6,10,28–34^. Behaviourally, sabotaging in danaine caterpillars changes over larval development. While first and second instars mainly trench, in older caterpillars the frequency of vein cutting increases. Isolating entire leaves from the latex flow by chewing a furrow into the petiole seems to be restricted to fifth- and sometimes fourth-instars of milkweed butterfly caterpillars ^3,6,19,28–31^.

**Figure 1.**
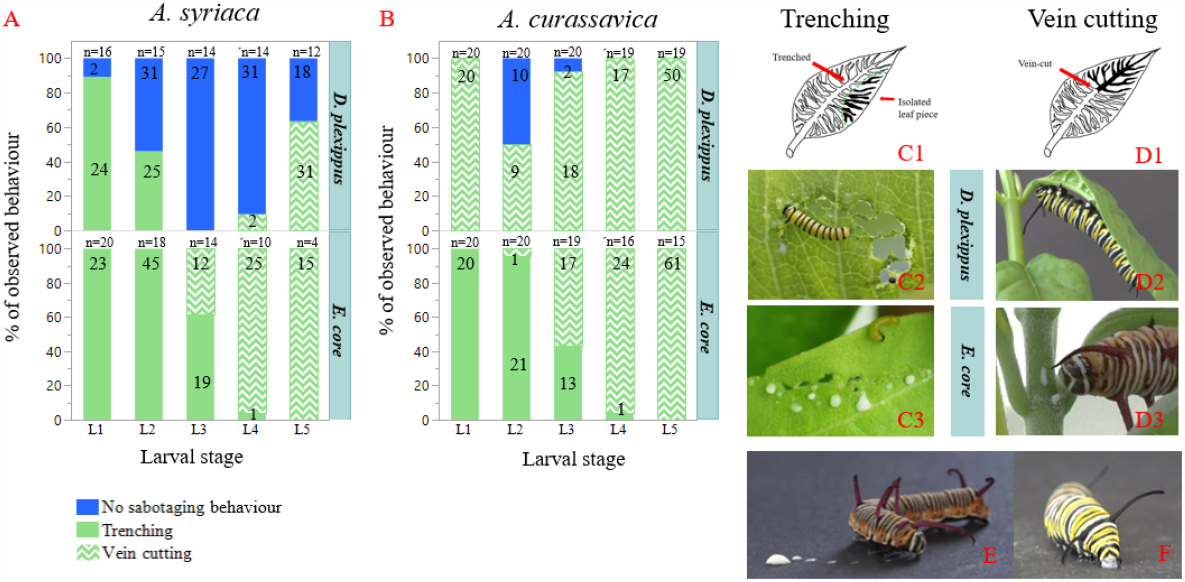
Sabotaging behaviour and latex drinking by caterpillars of the monarch butterfly (*D. plexippus*) and the common crow (*E. core*). Caterpillars (n = 20, per caterpillar species and plant species; total = 80) were raised singly on individual plants until pupation on *A. syriaca* (A) or on *A. curassavica* (B). Observed feeding behaviour was categorized into trenching (C1-3) (i.e. cutting a series of minor leaf veins, resulting in a trench), vein cutting (D1-3) (i.e. cutting a single major leaf vein) or no sabotaging behaviour (i.e. direct feeding on the leaf). Stacked bars represent mean percentages of sabotaging behaviour observations per instar, caterpillar species and hostplant species. Numbers in bars indicate total observations recorded. (F+E) Caterpillar behaviour after artificial latex feeding; *E. core* spitting out latex immediately (E); *D. plexippus* drinking latex (F).

When watching late instar *D. plexippus* caterpillars sabotaging *Asclepias curassavica* leaf petioles in the greenhouse, we never observed latex outflow and therefore hypothesized that caterpillars might drink exuding latex to acquire high amounts of cardenolides instead of avoiding it. Indeed, several authors have observed monarch caterpillars to “avidly imbibe” and to “actively seek and drink” milkweed latex ^35–38^ but the function and consistency of latex-drinking has never been addressed. Therefore, sabotaging behaviour in monarchs could represent co-option rather than the widely-assumed disarming of a famous plant defence ^3,6,8,12– 17,29,35,39–41^.

To address this question, we conducted experiments on *A. curassavica* and *Asclepias syriaca*, the two most important host plants of the monarch butterfly ^42 11^. For comparison, we used caterpillars of the closely related but non-sequestering common crow butterfly (*Euploea core*), which uses the same host plants and also shows similar sabotaging behaviour providing a suitable model system. First, we studied the frequency and type of sabotaging behaviour (i.e. trenching vs. vein cutting) in both butterfly species. Therefore, caterpillars individually raised on plants (initially n = 20 for each plant and insect species) were observed once a day and the following behaviours were recorded: ongoing sabotaging behaviour (trenching or vein cutting), caterpillar feeding with signs of preceding sabotaging behaviour, or caterpillar feeding without signs of preceding sabotage (see methods for details). For statistical analysis, behaviour was categorized as either sabotaging (trenching and vein cutting) or not-sabotaging.

While sabotaging preceded every feeding event of *E. core* on both milkweed species, *D. plexippus* caterpillars sabotaged milkweed inconsistently (Fig. 1A, B). Monarch caterpillars carried out sabotaging behaviour with varying consistency across larval stages (*F*_4,212.9_ = 12.44; *P <* 0.001) and host plant species (*F*_1,76.02_ = 27.3; *P <* 0.001). Especially on *A. syriaca*, sabotaging behaviour by monarch caterpillars became less frequent after the 1^st^ instar. While 1^st^ instar caterpillars trenched in 90% of observations, sabotaging behaviour in the 2^nd^ instar was only found in 46% of all observations, while being completely absent in caterpillars of the 3^rd^ instar (Fig 1A). On *A. curassavica, D. plexippus* sabotaging was more consistent (92 - 100% of observations in the 1^st^, 3^rd^, 4^th^, and 5^th^ instar) but also absent in 50% of 2^nd^ instar caterpillar observations (Fig. 1B). As carried out so inconsistently on natural host plants, our data suggest that sabotaging is not generally required by *D. plexippus* to avoid impairment by latex, especially in older caterpillars. In contrast, the consistent sabotaging behaviour observed in *E. core* clearly supports the hypothesis that sabotaging behaviour is a strategy to circumvent stickiness or toxicity of latex.

In a second experiment, we found a remarkable behavioural difference when we compared sabotaging behaviour between last instar caterpillars of *D. plexippus* and *E. core*. During petiole-cutting on both milkweed species, caterpillars of *E. core* always spit out latex multiple times close to the furrow and dabbed their mandibles on the plant stem (independent observations; n = 11 on *A. curassavica*; n = 2 on *A. syriaca*). When sabotaging individual petioles on *A. curassavica*, caterpillars removed latex nine times on average (± 3.39 SE, n = 7 independent observations). In contrast, latex-spitting was never observed in *D. plexippus* (independent observations; n = 12 on *A. curassavica*; n = 10 on *A. syriaca*; see the supplementary material for observations on sabotaging behaviour of both species, Fig. S1, S2; video S1, S2). In accordance, caterpillars of *E. core* took over six times longer to cut the petiole on *A. curassavica* compared to *D. plexippus* (see supplementary material for details).

To study the differences in sabotaging behaviour between the two caterpillar species in more detail, we applied *A. curassavica* latex with a pipette to the mouthparts of last instar caterpillars of both species (i.e. a single application of 6 μl per caterpillar, n = 12 for each species). In agreement with our previous observations on whole plants, ten out of twelve caterpillars of *E. core* spit out latex immediately (Fig. 1E). In contrast, all caterpillars of *D*.

*plexippus* (n = 12) imbibed latex instantly (*Fisher’s exact test, P* < 0.001, response variable “latex drinking: yes/no”, Fig. 1F; see supplemental video S3). Collectively, our observations support the notion that last instar caterpillars of *D. plexippus* drink latex, while *E. core* avoids and actively removes latex.

To test the hypothesis that monarch caterpillars may actively drink latex during petiole cutting, we conducted the following experiments. First, we evaluated the efficiency of latex-removal by monarch caterpillars during petiole cutting on *A. curassavica*. After removing the distal third of a leaf with a razor blade, we compared latex outflow of intact and petiole-cut *A. curassavica* leaves and found that petiole cutting reduced latex outflow substantially (*t*_17_ (equal variances) = -13.22, *P* < 0.001). Next, we tested ingestion of latex and cardenolides via chemical analysis of caterpillar regurgitations (i.e. foregut contents, Fig. S3). In support of our hypothesis, that monarchs drink while *E. core* caterpillars avoid latex, foregut contents of fifth instar monarch caterpillars (n = 10) harvested right after petiole cutting on *A. curassavica* had 10-fold higher cardenolide concentrations compared to caterpillars of *E. core* (n = 8) (*Wilcoxon, T =* 36, *z = - 3*.*51, P <* 0.001, Fig. 2A). Furthermore, monarch regurgitates harvested after petiole cutting had significantly higher cardenolide amounts than regurgitates collected before petiole cutting on both host plants (*A. curassavica*: *Wilcoxon T =* 55, *z =* - 3.748, *P <* 0.001, *A. syriaca Wilcoxon T* = 79, *z =* -1.992, *P =* 0.046, Fig. 2B) indicating latex drinking in monarchs.

**Figure 2.**
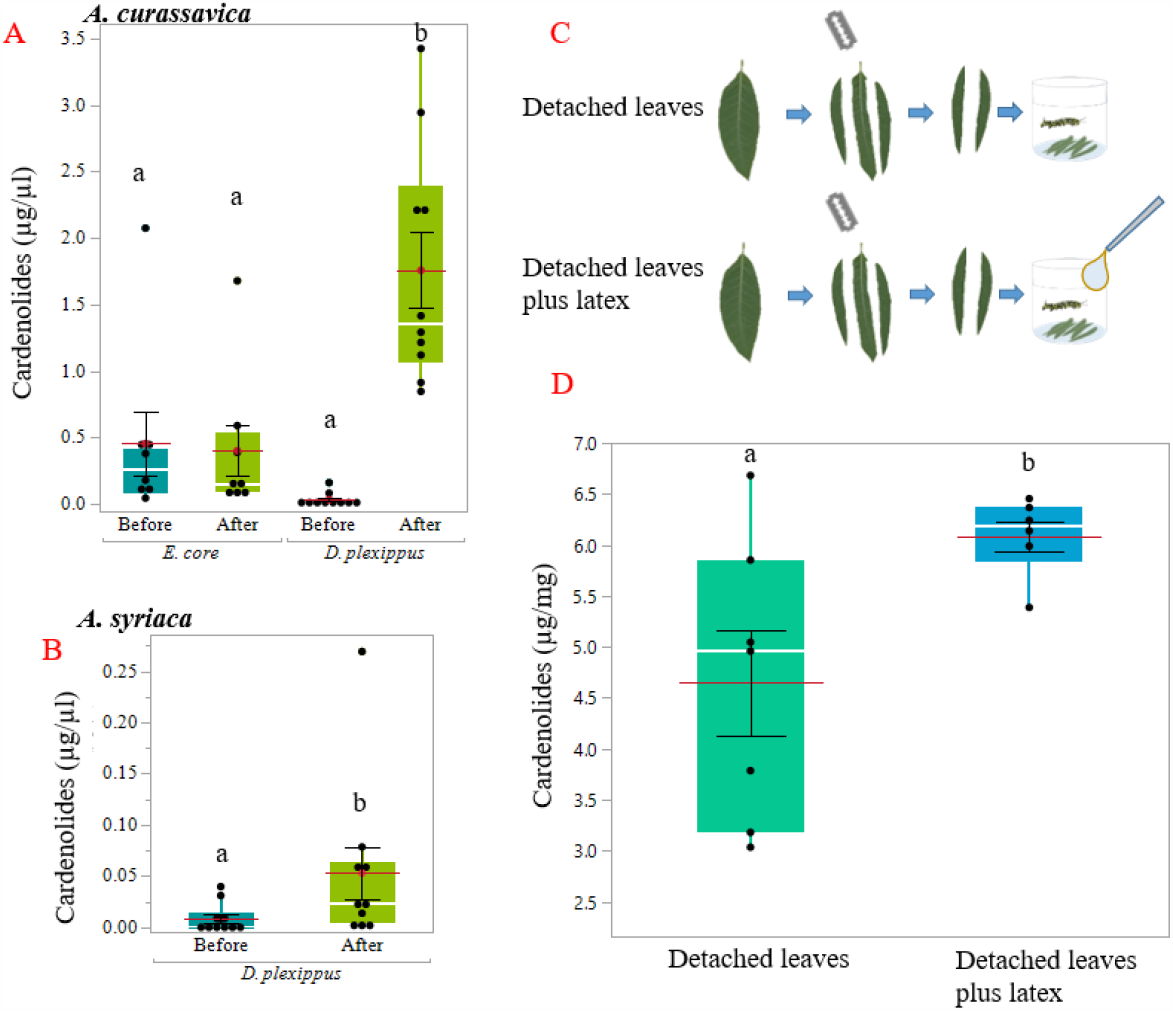
Cardenolide concentrations in foregut contents before and after sabotaging, sabotaged plants, and insect tissues. (A) Cardenolide concentration in foregut content of *D. plexippus* and *E. core* 5^th^ instar caterpillars before and after vein cutting on *A. curassavica*. Different letters above bars indicate significant differences between treatments. Boxes indicate interquartile ranges and white lines show medians. Black dots designate individual data points. Red lines represent means and black lines indicate the SE. (B) Cardenolide concentration in foregut content of *D. plexippus* before and after vein cutting on *A. syriaca*. (C) Experimental setup: caterpillars were raised from egg until pupation on latex-free (midrib removed) detached leaves of *A. curassavica* (top) or on the same diet but hand-fed with latex during the last instar (3 × 6 μl per day). (D) Concentration of sequestered cardenolides in adult butterflies between experimental diets.

Indeed, regurgitate samples might only represent a fraction of the total foregut content volume and the actual uptake of cardenolides might have been even higher. Based on our observations, a last-instar monarch caterpillar carries out petiole cutting at least three times per day and therefore the total amounts of cardenolides acquired by latex drinking from *A. curassavica* probably range around several hundreds of micrograms (total amounts of monarch foregut content samples range between 8.46 and 68.77 μg; 32.23 mean, ± 6.1 SE). Differences between cardenolide concentrations in regurgitates obtained from monarch caterpillar petiole-cutting on *A. syriaca* were less pronounced which is most likely due to the substantially lower cardenolide content of *A. syriaca* compared to *A. curassavica* (including latex) ^20^. Moreover, monarch caterpillars potentially obtain higher volumes of latex on *A. syriaca* plants in the field which are much larger compared to the pot-grown greenhouse plants used here.

We hypothesized that drinking latex in late instars maximizes cardenolide acquisition for transfer to adult butterflies. To investigate if latex drinking by caterpillars increases the amount of sequestered cardenolides in adult butterflies, we compared the cardenolide content of butterflies when reared individually on detached *A. curassavica* leaves (midrib of leaves removed to minimize latex content, n = 7) with butterflies raised on detached leaves when latex was supplied via a pipet to mimic drinking (n = 6, Fig. 2C). In support of our hypothesis, latex supply increased cardenolide concentrations in monarch butterflies (t_7_ = -2.66, *P* = 0.033, Fig. 2D; total cardenolide amounts of butterflies differed only marginally; t_11_ = -2.08, *P* = 0.062).

Previous studies demonstrated that latex exposure causes growth reduction or increases mortality of early instar monarch caterpillars ^19,29,43^. In line with these findings, we found that mortality of first instar caterpillars raised on detached *A. syriaca* leaves was significantly reduced compared to caterpillars grown on intact *A. syriaca* plants (*t*_135_ = -2.52, *P* = 0.039, Fig. 3A, see supplemental material for other stages, Fig. S4). Mortality of first instar caterpillars, however, did not differ when raised on intact or detached *A. curassavica* leaves (*t*_*129*_ *=* 0.01, *P* = 1, Fig. 3A, see supplemental material for other stages, Fig. S4). Based on these findings and our observations of latex consumption in late instars, we predicted a shift from avoidance behaviour to drinking over the course of larval ontogeny. To test this hypothesis, we raised monarch caterpillars on latex-free leaves of both plant species and applied latex of the respective host plant or water as a control on their mouthparts daily (volume of latex or water adjusted to caterpillar size, see methods for details).

**Figure 3.**
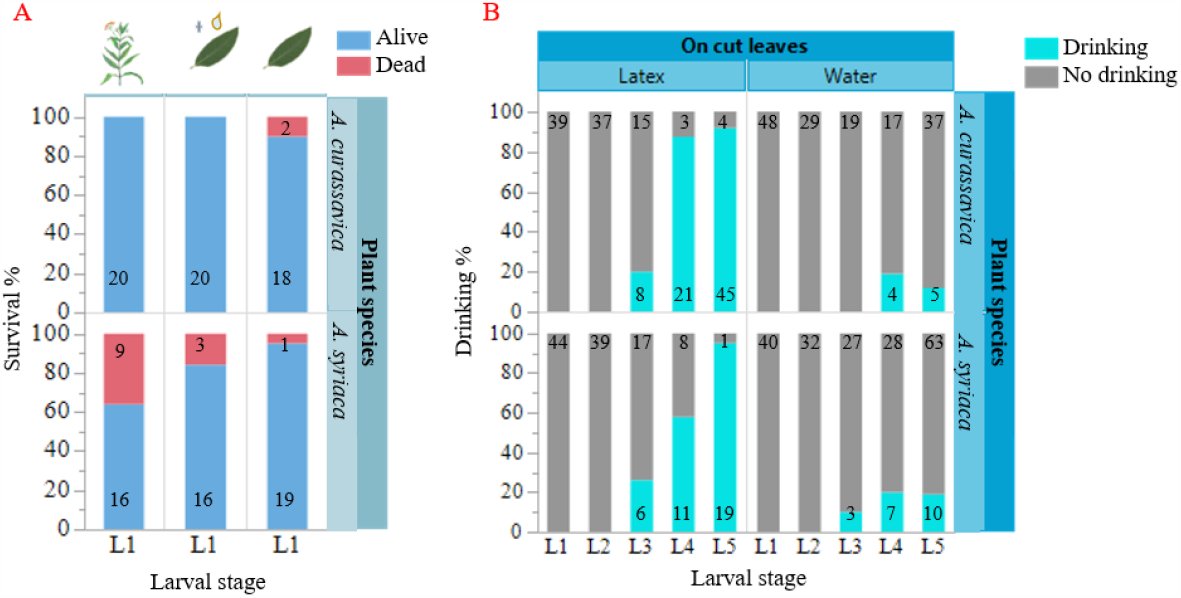
Young caterpillar survival on milkweed and latex drinking across caterpillar development. (A) survival of first instar monarch caterpillars raised on intact plants (left bars), detached leaves with daily application of 1 μl latex (centre bars), or detached leaves (right bars) of *A. curassavica* (top) and *A. syriaca* (bottom). Numbers in bars indicate the number of caterpillars observed. Data on caterpillar survival on intact plants were collected during the experiment shown in Fig. 1. (B) Proportion of caterpillars drinking latex of *A. curassavica* or *A. syriaca* (or water as a control) when offered manually across larval development. Caterpillars were raised on detached leaves of the respective milkweed species from which latex was obtained. Initially, n = 20 caterpillars per plant and treatment (i.e. 80 caterpillars in total) were observed. Caterpillars were offered latex or water once a day. Numbers in bars indicate the number of observed events per developmental instar.

As predicted, larval stage and type of liquid applied (i.e. latex or water), influenced the probability of caterpillar drinking, but was not affected by host plant species (*F*_4,47.79_ = 6.61, *P* < 0.001, and *F*_1,226.3_ = 136.77, *P* < 0.001 and F_1,221.5_ = 0.02, *P* = 0.898 respectively; Fig. 3B, see supplemental videos S4, S5). While younger caterpillars avoided latex, monarch caterpillars in the fifth instar were almost five times more likely to drink latex compared to water.

While sabotaging laticifers probably represents an ancestral trait of danaine caterpillars to overcome the stickiness and toxicity of milkweed latex, vein cutting in late instar monarch caterpillars most likely facilitates sequestration of cardenolides for defence against predators. We show that instead of avoiding it, older monarch caterpillars drink milkweed latex and incorporate copious amounts of cardenolide toxins leading to an increased net uptake into their body tissues. Caterpillar behaviour changes across larval ontogeny from latex avoidance in early instars to eager latex drinking in late instars. We suggest that an ancestral behaviour of tolerance was converted into a defence-related trait.

## Material and methods

### Cultivation of plants

To facilitate seed germination, *Asclepias syriaca* and *A. incarnata* seeds were nicked with a knife. We did not nick *A. curassavica* seeds. Seeds were then embedded in moist tissue, placed in Petri dishes, sealed with breathable tape, and stored at 28°C for 4 days in the dark. We planted seedlings in growing trays and transplanted them two weeks later into 11 × 11 × 12 cm (length x width x height) plastic pots (substrate 5, Klasmann-Deilmann GmbH mixed 3: 1 with sand). Starting one week after transplanting, plants were fertilized weekly with a 0.2 % dilution (v/v) of universal fertilizer (Wuxal Super N:P:K 8:8:6, Hauert MANNA Düngerwerke GmBH, Nürnberg, Germany). The bottom third of *A. curassavica* and *A. incarnata* pots was constantly immersed in water, while pots with *A. syriaca* were let to drain after watering. All plants were grown in a greenhouse at 23-28/ 21-24°C, with additional sources of artificial light. All experimental plants were around three months old. Seeds were originally obtained commercially.

### Butterfly rearing

*D. plexippus* (origin Portugal, stock provided by a local breeder) and *E. core* (origin Southeast Asia) were reared under greenhouse conditions (see above). Caterpillars of *D. plexippus* used for experiments were from a long-term colony maintained over multiple generations, while caterpillars of *E. cor*e either were offspring from freshly obtained adult specimens which were purchased commercially or were first-or second-generation offspring from commercially obtain specimens. We kept adult butterflies in 2 × 1 × 2 m flight cages and supplied them with a 10% sucrose solution and nectar plants (*A. curassavica, Lantana camara, Pentas lanceolata*). In addition, *Euploea core* was provided with chopped seeds of *Heliotropium indicum and H. foertherianum* ^45^. Butterflies were sprayed with water 8 times a day for 2 min using a fogging system (Micro Rain Systems, Altenburg, Germany) to increase humidity. For maintenance of butterfly colonies, caterpillars of both species were raised on potted *A. curassavica* plants under greenhouse conditions. For experiments, caterpillars were either reared on *A. curassavica, A. incarnata*, or *A. syriaca* according to the experimental requirements (see below).

### Experiments

#### (A) Sabotaging behaviour over ontogeny

We placed one egg each (n = 40) on the upper third of ca. three months old *A. curassavica* (n = 20) and *A. syriaca* (n = 20) plants to observe sabotaging behaviour of *D. plexippus* across larval development under ambient conditions in the greenhouse (see above). Sabotaging behaviour of caterpillars on both plant species that was either observed directly or was associated with a currently occupied caterpillar feeding site was recorded once a day from hatching to pupation. If caterpillars had cut more than one leaf vein in a circular manner, sabotaging was classified as trenching behaviour. Cutting a single furrow along the main leaf vein before feeding from the leaf tip was classified as vein cutting behaviour. Vein cutting was furthermore subdivided into three categories depending on where the main leaf vein was sabotaged (leaf petiole, leaf base, centre, or tip). It was not verified that caterpillars actually started feeding after observing ongoing sabotaging behaviour, but we never observed sabotaging events without subsequent feeding. Observations of direct feeding without any signs of associated sabotaging behaviour were categorized as “no sabotaging”. A similar experiment with *E. core* was conducted in climate chambers (Fitotron® HGC 1014, Weiss Technik, Reiskirchen, Germany) under increased humidity (27/24°C, 16/8h light/ dark, 80/ 50% humidity) to prevent drying out of the less robust early instar caterpillars of *E. core*. Again, the sample size was n = 40 eggs, but due to space limits in the climate chamber, two eggs were placed on one plant, resulting in n = 10 *A. syriaca* plants and n = 10 *A. curassavica* plants being sampled. Sabotaging behaviour was recorded once a day and classified as described for *D. plexippus* caterpillars. For statistical evaluation, the response data collected in this experiment was transformed to be binomial. For this, all categories of sabotaging behaviour (trenching or vein cutting on various positions of the leaf) was assigned the value “1”. The value “0” was assigned if no sabotaging behaviour was observed.

#### (B) Artificial latex feeding

We hand-fed a set of *A. curassavica* raised fifth instar caterpillars (n = 12) of both caterpillar species with 6 μl of *A. curassavica* latex under laboratory conditions and videotaped their reaction (Olympus OM-D E-M1 Mark III). Hand-feeding was done by holding the caterpillar with one hand and pipetting latex to the mandibles with the other hand (see video S3). A volume of 6 μl latex was found to be representative when petiole cutting was mimicked with a razor blade on additional *A. curassavica* plants (twice per plant; i.e. two averaged measurements per plant n = 10) and outflowing latex was measured with glass capillaries (6.4 ± 0.5 μl).

#### (C) Efficiency of latex-removal

*A. curassavica*-raised individual fifth-instar monarch caterpillars (n = 9) were placed on three-month-old *A. curassavica* plants. After petiole cutting, caterpillars were removed and the corresponding leaf was cut 5 cm from the tip with a razor blade. Outflowing latex was collected with a pre-weighed filter paper from both leaf portions (leaf base and tip) and weighed immediately. For comparison, we used leaves of the same size and age from intact *A. curassavica* plants of the same batch (n = 10) and collected latex as described above.

#### (D) Latex drinking during vein cutting

We tested if fifth instar caterpillars of *D. plexippus* drink latex during petiole cutting via chemical analyses of cardenolides in caterpillar regurgitates. Caterpillars of *D. plexippus* were raised from eggs on *A. syriaca* (n = 10) and *A. incarnata* (n = 10) plants under greenhouse conditions. After reaching the fifth instar, we induced regurgitation in actively feeding caterpillars by tweaking the caterpillars’ abdomen with tweezers and collected the full amount of regurgitate with glass capillaries (blank). In addition, we recorded sample volumes since the amount varied across caterpillar individuals. Next, we transferred *A. syriaca*-reared caterpillars to *A. curassavica* plants and *A. incarnata*-reared caterpillars to *A. syriaca* plants (one caterpillar per plant, 3-months-old plants). Caterpillars were raised on *A. syriaca* and *A. incarnata* since the cardenolides produced by these milkweed species differ structurally from the cardenolides found in *A. curassavica* or *A. syriaca*, respectively. Consequently, cardenolides ingested during vein cutting can be easily differentiated from cardenolides present in the host plant. In addition, both, *A. syriaca* and *A. incarnata*, produce only comparatively low amounts of cardenolides^46^. All caterpillars started petiole cutting within 1 hour. When petiole cutting was finalized and caterpillars started walking to the leaf tip for feeding, a second regurgitate sample was collected as described above. All samples were extracted and the cardenolide content was quantified using HPLC (see below for methods of extraction and quantification). In parallel, we tested if caterpillars of *E. core* avoid oral uptake of latex during vein cutting in an according experiment. We used caterpillars raised from egg on *A. curassavica* (n = 8) since *E. core* did not grow well on *A. syriaca*. Before the experiment, fifth instar caterpillars were transferred to uninjured *A. curassavica* and samples were collected as described above. Due to the tendency of *E. core* to chew furrows above the petiole (six out of eight), we also included caterpillars which carried out vein cutting within the proximal third of the leaf.

#### (E) Latex drinking and butterfly toxicity

To evaluate the effect of latex drinking in caterpillars on overall toxicity of monarch butterflies, we raised caterpillars either (1) on detached *A. curassavica* leaves (see below for details), or (2) on detached leaves with additional *A. curassavica* latex-feeding of caterpillars. For both diets, caterpillars were raised from egg to pupae. Caterpillars were individually raised in plastic containers (125 ml until L1-L4, 500 ml L5, n = 7) lined with moist filter paper and supplied with freshly detached *A. curassavica* leaves after excision of the midrib to maximize latex removal. A maximum of two leaves per plant per day were collected and leaves were removed with a razor blade to avoid cardenolide induction in donor plants by harvesting^47^. For (2), caterpillars were treated as described for (1) but were additionally hand-fed with 6 μl of *A. curassavica* latex three times per day using a pipette from the fifth instar onwards (n = 6). Supplementation with latex was limited to the 5^th^ instar since caterpillars showed a marked increase in the frequency of petiole cutting during the last instar (see Fig. S2). As mentioned above, 6 μl was found to be a typical volume of latex-outflow and three petiole cutting events per day represent a conservative estimate of daily petiole cutting frequency in 5^th^ instar monarch caterpillars. Representative leaves without midrib (n = 7) and native leaves collected adjacently to leaves used for the experiment (n = 7) had similar concentrations of cardenolides (F_2,22_=0.03, p=0.97). After pupation, we transferred pupae to room temperature for hatching. After wing hardening, butterflies were frozen at -80°C. All samples (butterflies and leaves) were stored at -80°C, freeze-dried (Alpha 2-4 LDplus, Martin Christ, Osterode, Germany), weighed on a microbalance (0.001 g; Cubis II, Sartorius Corporate Administration GmbH, Göttingen, Germany), extracted, and analysed via HPLC-DAD as described below. Unequal sample sizes in this experiment are due to high mortality of caterpillars reared in containers probably due to adverse environmental conditions (high temperatures) in the greenhouse.

#### (F) Latex influence on larval instars

We estimated the effect of plant latex on caterpillar survival by rearing caterpillar on detached leaves, with and without additional latex supply. Caterpillars of *D. plexippus* were raised from egg in plastic containers lined with moist filter paper (cups of 125 ml until L3, 500 ml from L4 on) on detached leaves of *A. syriaca* or *A. curassavica* (27/24°C, 16/8h light/ dark, 60/0% humidity in a Fitotron® HGC 1014 chamber, Weiss Technik, Reiskirchen, Germany). On both plant species, half of the caterpillars (initially n = 20) were fed with latex of the corresponding plant once a day (L1-L2 = 1 μl, L3 = 2 μl, L4 = 4 μl, L5 = 6 μl). The other half (initially n = 20) was fed with an according volume of water. Latex or water feeding was carried out with a pipette. The behaviour upon latex or water exposition and survival of caterpillars was recorded across caterpillar development. Survival of caterpillars on detached leaves was compared to survival of caterpillars raised on intact plants during experiment (A).

### Extraction of foregut contents

We collected samples in 2 ml screw cap micro tubes containing 0.9 g of zirconia beads (2.3 mm, Carl Roth GmbH, Karlsruhe, Germany) and 1 ml methanol. Samples were homogenized twice with a FastPrep homogenizer (MP Biomedicals, Eschwege, Germany) for 45 sec (6.5 m/s). After 3 min of centrifugation (16,000 g at 22°C; 5417R, Eppendorf, Hamburg, Germany), we transferred the supernatant into new 2 ml screw cap micro tubes. This procedure was repeated twice. Subsequently, samples were evaporated to dryness in a vacuum centrifuge (RVC 2-25 CDplus, Martin Christ, Osterode, Germany). Finally, dried residues were dissolved in 100 μl methanol, agitated in the FastPrep instrument (45 sec; 6.5 m/s) and centrifuged (3 min, 16,000 g, 22°C). Subsequently, supernatants were filtered into HPLC vials using Rotilabo ® syringe filters (nylon, 0.45 μm pore size, Ø 13 mm, Carl Roth GmbH & Co.KG, Karlsruhe, Germany).

### Extraction of butterflies and leaves

Freeze-dried samples were transferred into a 15 ml centrifuge tube containing two ceramic beads (6.35 mm, MP Biomedicals, Graffenstaden, France) and 2 ml of methanol. The samples were homogenized in the FastPrep instrument as described above. After centrifugation for 10 min (1000 g at 22°C; Labofuge 400R, Heraeus Instruments, Osterode, Germany), we transferred 2 ml supernatants into a glass test tube. Extraction of samples was repeated once more with 2 ml methanol. Pooled supernatants (4 ml) were evaporated to dryness in a vacuum centrifuge (RVC 2-25 CDplus, Martin Christ, Osterode, Germany). Dried residues were dissolved in 500 μl methanol and transferred into a 2 ml screw cap micro tube. This procedure was repeated twice using an ultrasonic bath during the last round to completely dissolve any residue. The 1.5 ml of sample was evaporated again to dryness in the vacuum centrifuge. Samples were dissolved in 200 μl methanol (butterflies in 500 μl), homogenized (45 sec; 6.5 m/s), centrifuged (3 min, 16,000 g, 22°C), and filtered (nylon filter 0.45 μm, Ø 13 mm) into HPLC vials.

### Cardenolide quantification

We quantified the cardenolide content of the samples using an Agilent Infinity 1260 II HPLC system (Agilent Technologies, USA) using an EC 150/4.6 NUCLEODUR C18 Gravity column (3 μm particle size, 150 mm × 4.6 mm, Macherey-Nagel, Düren, Germany). We injected 15 μl of sample which was eluted at a constant flow rate of 0.7 ml/min using a water/acetonitrile gradient as follows: 0-25 min 16% acetonitrile; 25 min 70% acetonitrile; 30-35 min 95% acetonitrile. Peaks were detected at 218 ± 4 nm using a diode array detector (DAD). Absorbance spectra were recorded between 200-400 nm. Peaks with a characteristic symmetric absorption spectrum with a maximum between 218 and 222 nm were classified as cardenolides ^48^. Cardenolides were quantified at 218 ± 4 nm using a digitoxin calibration curve (5, 10, 25 50, 100, 250, 500, 750, and 1000 μg/ml). All cardenolide peaks in one sample were summed up to calculate total cardenolide concentrations.

### Data analysis

Statistical analysis was performed with SAS 9.4 software (SAS Institute, Cary, NC). For statistical codes and outputs please see the statistical documentation. P-values < 0.05 were considered statistically significant. Data collected in this experiment were either metric or binomial (see below).

Metric data (datasets on trials C, D, E, G (in supplemental material)), were tested for normal distribution prior to analysis (Shapiro-Wilk test). Normally distributed metric datasets were analysed via *t-test* (C, E). We used the *t-test* of Satterthwaite for detection of unequal variances. If normality was rejected (D, G), analysis was carried out with Wilcoxon rank sum test for two samples. Binomial data were analysed using generalized linear mixed (GLMM) models (A, F). For these trials, caterpillars were tested multiple times throughout their larval development, therefore a repeated effect was fitted with the subject level individual caterpillar. The assumed structure of correlation (covariance structure) was autoregressive (1). For the data on survival (included in F) no covariance structure was assumed as the assessed event (death) can only occur once. Overdispersion of the generalized linear mixed model was rejected if Generalized Chisq. / DF= < 1. For dataset (B) a Fisher’s exact test was used. For all analyses, no outliers were excluded. The graphical representation of our data was generated with JMP Pro 16.1.

## Acknowledgements

We thank Margit Schmidtke and Sarah Rißmann for help with butterfly rearing, plant growing, and technical support. Johanna Weber, Hermann Falkenhahn, Sabrina Stiehler, and Martin Planke supported the early stages of this work. Jens Hartung provided valuable statistical support. This manuscript was improved by discussions with Anurag Agrawal and Tobias Züst.

## Author contributions

Conceptualization: GP, AB

Methodology: AB, GP

Investigation: AB

Data analysis: RB, AB, GP

Visualization: AB, GP, RB

Funding acquisition: GP

Project administration: GP

Supervision: GP

Writing – original draft: AB, GP

Writing – review & editing: AB, GP, RB

## Data and materials availability

All data underlying our study will be deposited at the dryad data repository (https://datadryad.org/stash) in due course and will be available upon acceptance of the manuscript. No restrictions on data availability apply.

## Supplementary Materials

SI Guide

Supplementary information

Figs. S1 to S4

Movies S1 to S5

