## Supplementary Materials for "Monarchs sabotage milkweed to acquire toxins, not to disarm plant defence"

**Supplementary Material**

**Methods**

**(G) Petiole cutting duration of last instars**

We measured the duration of vein cutting at the leaf petiole by last instar caterpillars on *A. curassavica* (*E. core*: n = 11, *D.* plexippus: n = 12) and *A. syriaca (E. core*: n = 2, *D. plexippus*: n = 10) and videotaped a subset of caterpillars (Olympus OM-D E-M1 Mark III). Specifically, we assessed the time caterpillars needed to furrow the leaf petiole until they turned around to start feeding from the leaf tip. Five out of eleven caterpillars of *E. core* on *A. curassavica* sabotaged at the midrib on the leaf blade, while *D. plexippus* exclusively cut furrows into leaf petioles. The experiment was carried out under laboratory conditions for *D. plexippus* and in the greenhouse or climate chamber for *E. core* (conditions as described above). We note, that we did not observe differences of petiole cutting duration and behaviour, when observing additional caterpillars of *D. plexippus* under greenhouse conditions, therefore potential environmental differences did not seem to influence the duration of sabotaging behaviour.

**Results**

On *A. curassavica*, caterpillars of *E. core* spent on average 19.63 min (± 2.24 SE, n = 11) for petiole cutting while caterpillars of *D. plexippus* took only 3.16 min (± 0.34 SE, n = 12, *Wilcoxon*, $T = 198$, $z = 4.032$ $P < 0.001$). On *A. syriaca*, petiole cutting took much longer for both caterpillar species but the relative differences remained (*D. plexippus*: 28.04 min ± 3.19, n = 10, *E. core*: 74.73 min ± 8.64, n = 2, (means ± SE, no statistical analysis due to low sample size).

### Figures

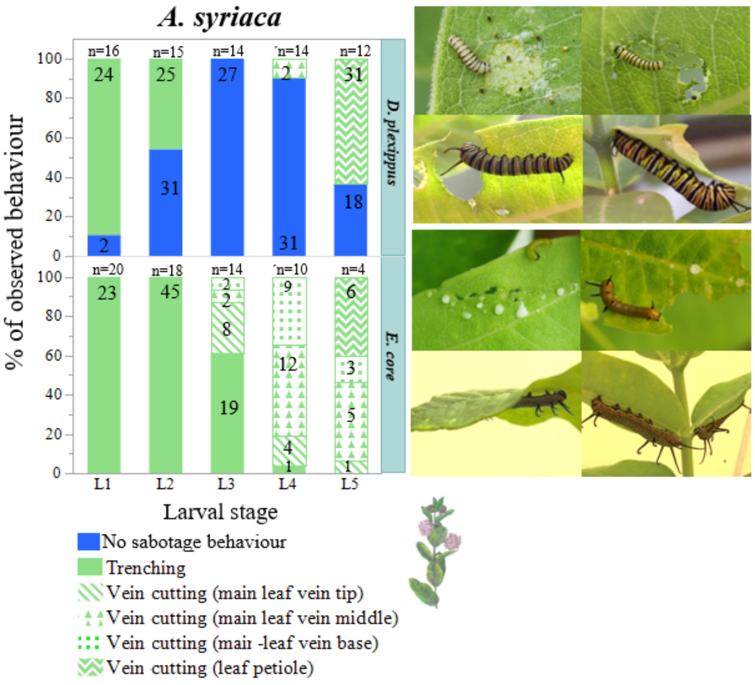


**Fig. S1.** Different types of sabotaging behaviour across caterpillar development on *A. syriaca*. Sabotaging behaviour was recorded daily in both, caterpillars of *D. plexippus* and *E. core* (left). Mean percentages of sabotaging behaviour type per instar are shown (*D. plexippus* top; *E. core* bottom). Numbers in bars indicate the total of recorded events per developmental instar. Images (right) illustrate representative behaviours observed in L1, L2, L3 and L5 caterpillars of both species (*D. plexippus* top; *E. core* bottom).

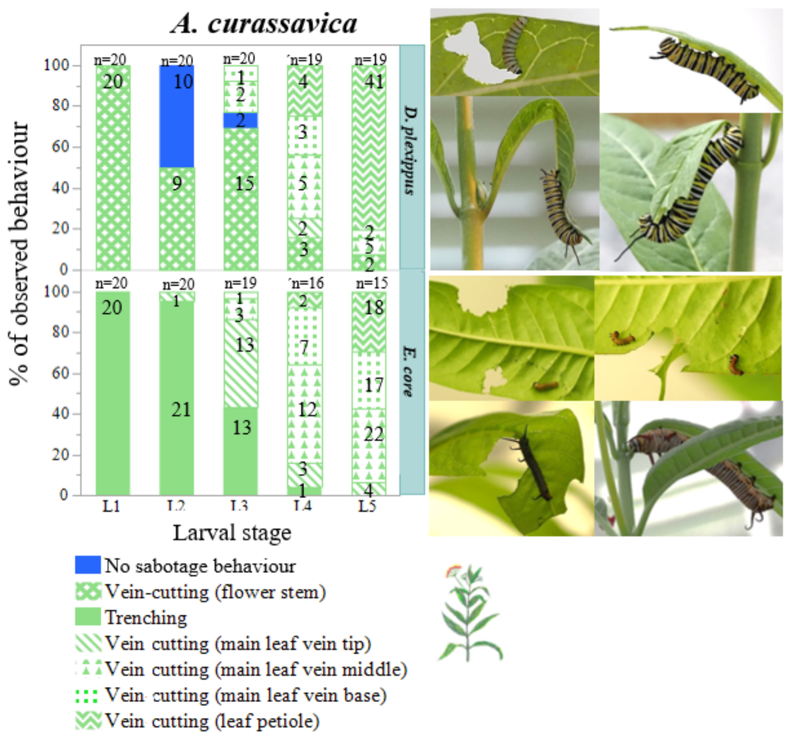


**Fig. S2.** Different types of sabotaging behaviour across caterpillar development on *A. curassavica*. For details see legend of figure S1.

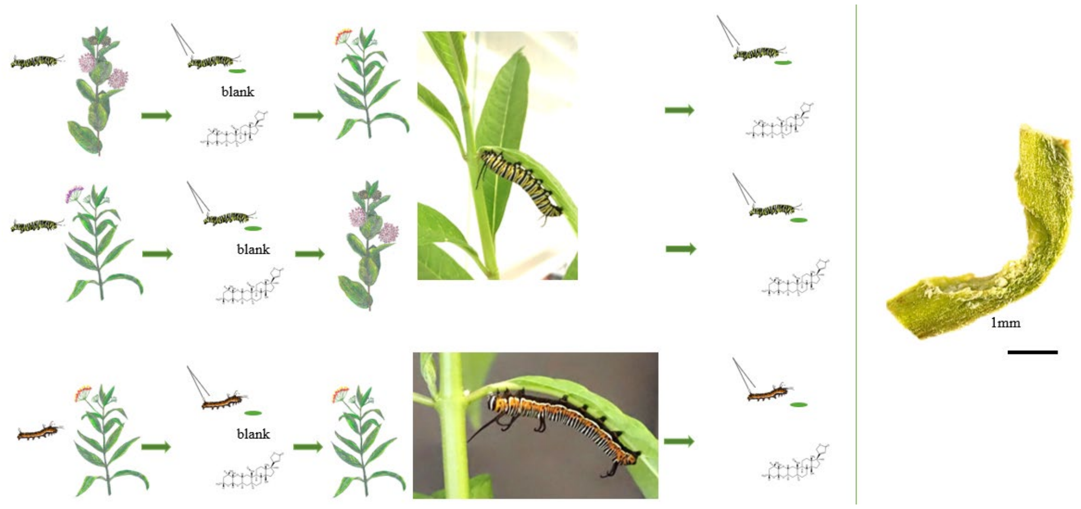

**Fig. S3.** Experimental setup for measuring cardenolide concentrations in foregut contents of *D. plexippus* and *E. core* before and after vein cutting on *A. curassavica*. *D. plexippus* caterpillars were raised on *A. syriaca* for measuring cardenolide concentrations after petiole cutting on *A. curassavica* and raised on *A. incarnata* for measuring cardenolide concentrations after petiole cutting on *A. syriaca*. Caterpillars of *E. core* were raised on *A. curassavica* and cardenolide concentrations after petiole cutting were measured on the same plant. The picture on the right illustrates a furrow created by *D. plexippus* caterpillar during petiole cutting.

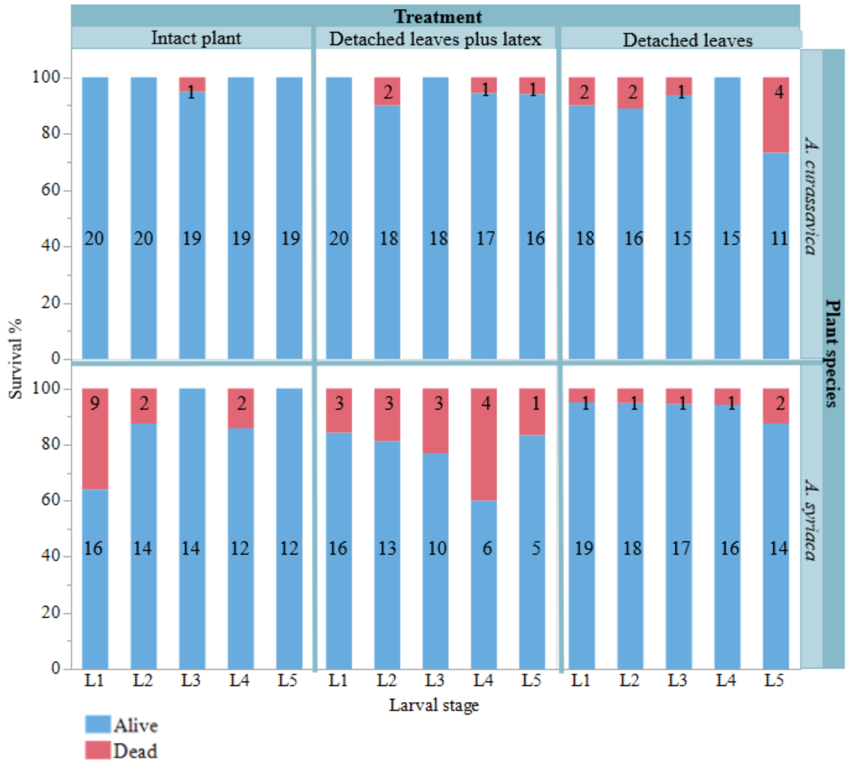


**Fig. S4** Survival of *D. plexippus* caterpillars raised on intact plants of either *A. curassavica* or *A. syriaca* plants or detached leaves with and without daily supply of latex via hand-feeding. For latex feeding, latex of the corresponding plant was used and the volume was increased over larval development (L1-L2 = 1 µl, L3 = 2 µl, L4 = 4 µl, L5 = 6 µl). Numbers in bars indicate the number of caterpillars observed.

### Movies

**Movie S1.**

Sabotage behaviour of fifth instar *D. plexippus* on *A. curassavica* and *A. syriaca*

**Movie S2.**

Sabotage behaviour of fifth instar *E. core* on *A. curassavica* and *A. syriaca*

**Movie S3.**

Latex acceptance through hand feeding of *E. core* and *D. plexippus*

**Movie S4.**

Latex drinking over larval stages of *D. plexippus* on detached *A. syriaca* leaves

**Movie S5.**

Latex drinking over larval stages of *D. plexippus* on detached *A. curassavica* leaves
